# The ParB-CTP Cycle Activates Phase Separation in Bacterial DNA Segregation

**DOI:** 10.1101/2025.02.17.638629

**Authors:** Linda Delimi, Perrine Revoil, Hicham Sekkouri Alaoui, Jérôme Rech, Jean-Yves Bouet, Jean-Charles Walter

## Abstract

Cell function relies on liquid-like membraneless organelles formed through phase transitions, yet the mechanisms ensuring their specificity and rapid assembly remain poorly understood. In bacterial chromosome segregation via the ParAB*S* system, hundreds of ParB proteins are recruited around the centromere-like *parS* sequence forming the partition complex. Recent studies have shown that ParB binds CTP and undergoes cycles of loading and unloading near *parS*, however, this accounts for the recruitment of only a small fraction of ParB molecules, leaving its role unclear. Separately, a lattice gas model with fixed interaction energy has been proposed to describe ParB phase separation, but it fails to explain key experimental observations, including the absence of droplets in ParB variants. We reconcile these two perspectives by proposing that the ParB-CTP cycle acts as a molecular switch that enhances ParB-ParB interactions, triggering phase transition from vapor to liquid-like condensates. Our hypothesis is supported by numerical simulations of droplet formation and experiments showing that ParB variants disrupting the CTP cycle fail to undergo phase separation. These findings establish a mechanistic framework for ParB-CTP-mediated phase transitions and may have broader implications for understanding the spatial control of intracellular condensate formation.

## Introduction

The formation of membraneless compartments with high concentrations of proteins, also called membrane-less organelles, relies on the physical mechanism of phase separation [1]. The resulting compartments consist of droplet-like objects that share liquid properties. It is an efficient way to quickly recruit, with a small amount of energy, a population of proteins to, for example, catalyze a chemical reaction. A prominent early example of phase separation is the assembly of P granules in *C. elegans* [2]. These condensates were among the first to show how weak, multivalent interactions facilitate the rapid and selective organization of biomolecules [1]. Similarly, in bacteria, several phase-separating systems exhibit liquid-like properties, such as in RNA processing regulation (BR-bodies) and the initiation of cell division (FtsZ-SlmA) [3]. Despite these insights, mechanisms leading to highly controlled phase transitions are still not well understood.

Bacterial DNA segregation offers a system for investigating the onset of droplet formation *in vivo*. Faithful DNA segregation is an essential biological process to ensure that each offspring receives at least one copy of each replicon, chromosome or plasmid. The ParAB*S* systems are the main active DNA partitioning systems used by bacterial chromosomes and most low-copynumber plasmids [4]. It consists of ParA, a Walker-type ATPase, and ParB, a CTPase that binds to *parS* centromere sites with high affinity. The initial step is the binding of a few ParB onto *parS* triggering the recruitment of hundreds of ParB proteins around *parS, i*.*e*., most of the intracellular ParB population (*>* 90%) [5]. These large nucleoprotein complexes are called partition complexes. ParA mediates their separation by activating its ATPase activity, which is catalyzed by the high local concentration of ParB [6, 7].

The partition complexes display several liquid-like properties *in vivo*, including: the averaged spherical shape of the condensate, which minimizes the surface tension, the ability of ParB foci to fuse, the intracellular mobility of ParB, which is ∼100 times slower within ParB complexes compared to the surrounding diluted phase, the rapid exchange of ParB between distinct partition complexes, occurring within a few minutes [8–10]. Furthermore, *in vitro* studies have demonstrated the self-organization of ParB into droplets [11, 12].

The lattice gas (LG) is a paradigmatic model for phase transitions where nearest-neighbour particles display attractive interaction. The LG exhibits two distinct phases: a vapor phase, where particles are randomly distributed within the volume, and a liquid-vapor (LV) coexistence phase, where the liquid coexists with a residual vapor phase. Within the LV phase, a metastable and a stable region can be identified. We previously investigated the partition complex formation, using this model at fixed ParB-ParB interaction energy in the metastable region. However, within this framework, the timescale of the complex formation is orders of magnitude larger than the cell cycle, making it biologically irrelevant [10].

New critical insight into this process is provided via ParB binding to CTP, and the subsequent ParB-CTP cycle, as recently described by the Clamping & Sliding (CS) mechanism [13]. The CS model involves a sequential multi-step process: (i) the specific binding of ParB to *parS*, a 16-bp DNA motif [14–16], (ii) the binding of CTP to *parS*-bound ParB [17, 18], followed by (iii) the conversion of ParB into a clamp and its ejection from *parS* due to steric clash upon ParB remodeling [18], (iv) the subsequent diffusion over *parS*-proximal DNA [18], and lastly (v) the unloading of ParB after the clamp reopens [19].

The CS model was proposed to explain the recruitment of most intracellular ParB to DNA [9, 20]. However, our findings reveal that CS alone can only account for a small fraction of ParB onto DNA (*<* 10 ParB proteins) and the observed loading and unloading timescales are inconsistent with experiments [10, 13]. New evidence from both *in vitro* experiments and simulations indicates that recruitment events can occur in *trans* but are generally sparse [20, 21]. Importantly, the binding of ParB to both *parS* and CTP is essential for complex formation. In the absence of *parS*, or when ParB variants lack CTP-binding ability, partition complexes fail to form [5, 9, 22, 23]. Additionally, the CTPase activity of ParB facilitates the release of clamped ParB from *parS*-proximal DNA [9, 11, 24]. Despite these findings, the functional role of CTP binding remains unclear and is still under debate.

To reconcile the LG and CS mechanisms, we hypothesize that the ParB-CTP cycle activates the phase transition by enhancing ParB-ParB interactions, providing a regulatory mechanism to fine-tune the formation and stability of partition complexes. Firstly, we outline this hypothesis and describe the theoretical basis of our model that couples LG framework with the CS mechanism. Secondly, we present kinetic simulations of partition complex formation, demonstrating that phase separation depends on an activation in order to occur in a biologically relevant timescale. Thirdly, we provide experimental evidence showing that modifications of the ParB-CTP cycle impair partition complex formation. We employed the well-characterized ParAB*S* system of plasmid F, hereafter called ParAB*S*_F_ [25]. Specifically, we engineered ParB_F_ variants in the CTP binding and hydrolysis motifs, both of which exhibited loss of the ParB_F_ droplet formation. Finally, we conclude by discussing the implications of these findings for understanding ParB-mediated DNA segregation.

## Material and Methods

### Monte Carlo simulations

The Monte Carlo (MC) simulations were performed using the Metropolis algorithm using the Hamiltonian of the LG to study the kinetics of ParB droplet formation [26]. At each MC iteration, the system attempts random particle movements towards an empty site of the lattice. The simple cubic lattice serves here as a mean field representation of DNA, considering plasmid and chromosomal DNA indistinctively. The lattice unit represents a binding site for ParB, which is approximately 16 bp of DNA and hard wall boundary conditions are implemented. The *parS* sites, which act as nucleation points for ParB droplets, are placed at the center of the lattice. The binding energy between ParB and *parS* (*ϵ*) is typically much larger than the binding energies of ParB-ParB or ParB-DNA interactions. This was characterized experimentally by high affinities (∼ 2 nM) compared to low affinities for ParB interactions with non-specific DNA (∼ 0.3–0.5mM) and ParB-ParB interactions (∼ 0.3–1 mM) [5, 6, 27]. To reflect this in our simulations, the binding site is fixed at *ϵ* = 10*kT*. The ParB_F_ diffusion coefficient (*D*), has been measured experimentally at approximately 1*µm*^2^*/s* [9,10] for freely diffusing ParB. Using this value, we calibrate the Monte Carlo iterations, with each MC step corresponding to one second of real time.

For equilibrium simulations presented in Fig.2(A), each change is followed by thermalization over 2 ×10^6^ MC steps, with subsequent sampling performed every 1000 steps. Additionally, we performed 30 independent cycles of increasing and decreasing of the interaction energy between ParB, *J*.

### Definition and analyses of the skewness

The skewness *S*, which quantifies the asymmetry in the pixel intensity distribution of average ⟨*I*⟩ and standard deviation *σ* is given by:

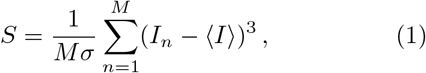

where *M* is the number of pixels in the image to analyze. Skewness measurements were performed using MicrobeJ softwares and the plugin “skewness”.

### Bacterial strains, plasmids and growth conditions

Strains and plasmids are listed in Supplemental Table S3. ParB_F_ variants in boxIII were constructed by Quik Change mutagenesis frompDAG114 (*parB* wt) or pJYB234(*parB* -mVenus). All constructs were verified by DNA sequencing. Cultures were essentially grown at 37°C with aeration in LB containing thymine (10 *µg*.*ml*^*−*1^) and chloramphenicol (10 *µg*.*ml*^*−*1^) as appropriate. For microscopy assay, cultures were grown at 30°C with aeration in MGC (M9 minimal medium supplemented with 0.4 % glucose, 0.2 % casamino acids, 1 mM MgSO4, 0.1 mM CaCl2, 1 *µg*.*ml*^*−*1^ thiamine, 20 *µg*.*ml*^*−*1^ leucine and 40 *µg*.*ml*^*−*1^ thymine).

### Plasmid stability assays

Stability of mini-F plasmids was assayed in strain DLT1215 grown over 25 generations in MGC at 30°C, and subsequently plated on LB agar medium, replica plating to medium with chloramphenicol, and calculating loss rates from the fractions of each sample resistant to chloramphenicol, as previously described [28].

### Epifluorescence microscopy

Exponentially growing cultures were deposited on slides coated with a 1% agarose buffered solution and imaged as previously described [29], using an Eclipse TI-E/B wide field epifluorescence microscope. Snapshots were taken using a phase contrast objective (CFI Plan Fluor DLL 100X oil NA1.3) and Semrock filters sets for YFP (Ex: 500BP24; DM: 520; Em: 542BP27) and Cy3 (Ex:531BP40; DM: 562; Em: 593BP40) with an exposure time range of 0.1–0.5 s. Nis-Elements AR software (Nikon) was used for image capture and editing. Image analyses were performed using ImageJ and MicrobeJ softwares.

### Chromatin immunoprecipitation DNA sequencing(ChIP-seq)

High resolution ChIP-sequencing was carried out using an affinity-purified anti-ParB antibody as previously described [30,31]. The ChIP-seq data (each assays with relevant information are summarized in Table S3) were processed using RStudio software with custom scripts (available upon request). ParB reads were counted at the center of the DNA fragments, taking into account the average size of each DNA library. Background levels on the plasmid F were determined by averaging ParB reads within genomic positions ranging from 1-to 31-Kbp and 74-to 99-Kbp. Following background subtraction, the ParB reads were binned every 20-bp, by applying a smoothing function with averaging over a 40-bp window with a step size of 20-bp. ParB density was obtained by normalizing reads relative to the highest value. For quantitative comparison between input and IP, the number of reads were normalized relative to the number of mapped reads. The raw and processed ChIP-seq data reported in this paper have been deposited in Gene Expression Omnibus database (GEO, NCBI) and are accessible through GEO Series accession numbers GSE256357 (ParB_F_) and GSE289093 (ParB_F_-N153A and ParB_F_-N156A). Unpublished custom codes are available upon request.

## Results

### Lattice Gas coupled to CTP activation of ParB

In this section, we present a two-step model combining a phase separation and a CTP activation during the CS cycle. We show a schematic representation of the ParB-DNA complex in Fig. 1A, showcasing the macroscopic behaviors of ParB observed experimentally. We define by *J* the interaction energy between ParB. On top of the panel, in the absence of *parS* or for ParB variants displaying modification of the ParB-CTP cycle, no droplets are observed [5, 9, 11, 22, 23]. ParB are then represented by green dots and feature an interaction energy *J*_V_. The bottom of the panel depicts the wild type (WT) scenario, where activated ParBs (magenta dots, interaction energy *J*_LV_) form droplets around *parS* sites. In the latter case, a majority of ParB proteins (∼ 95%) forms a dense complex in equilibrium with a small fraction (∼ 5%) outside [10]. These observations led to the development of the LG model to account for those macroscopic behaviors, thus enhancing the role of molecular ParB interactions.

**Figure 1.**
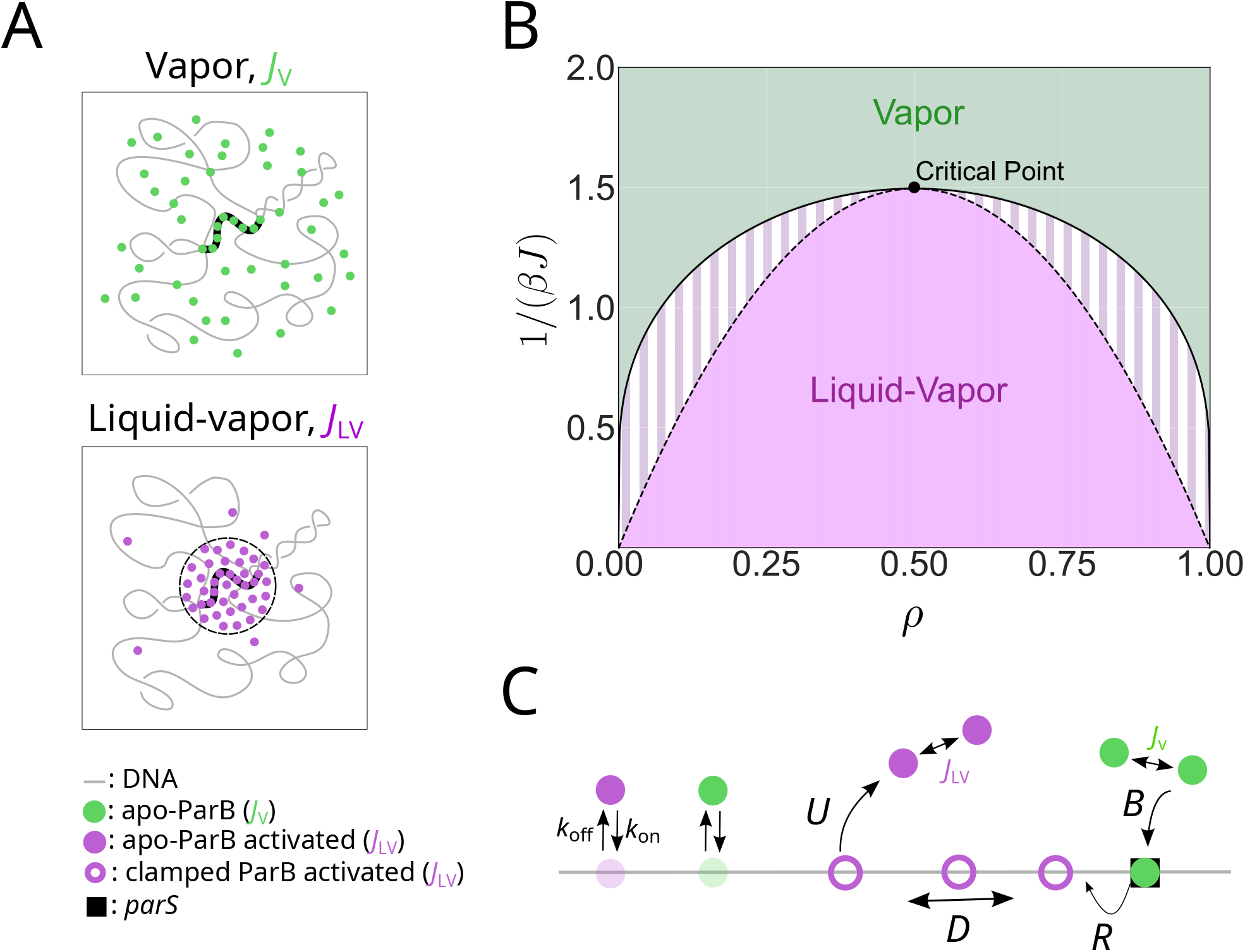
Modeling of phase transition controlled by ParB-CTP cycle activation. **A** Illustration of the ParB-DNA complex macroscopic behaviors. (top) Mutants with ParB variants disrupting ParB-CTP cycle fail to form complexes (ParB as green dots, with interacting energy *J*_V_). (bottom) WT cells display liquid-like ParB droplets around *parS*, with activated ParB (magenta dot, with interacting energy *J*_LV_). **B** Phase diagram of a LG in the (1*/*(*βJ*), *ρ*) plane (mean field solution [32]). The coexistence line (solid line) separates the vapor (green) from the liquid-vapor (purple) phase. The spinodal line (dashed line) demarcates the metastable region (white and magenta stripes) within the LV phase. **C** Schematic representation of ParB activation in the Clamping & Sliding stage. Apo-ParB specifically binds to *parS* (black square) at the rate *B*, which increases ParB affinity for CTP. CTP-binding induces a conformational change converting ParB-CTP into a clamp (magenta circle) over the DNA, leading to its ejection from *parS* at the rate *R*. For simplicity we hypothesize that the activation takes place between this latter step and the unloading from DNA. Clamped ParB-CTP can diffuse along the DNA, with a diffusion coefficient *D*, until being unloaded from non-specific-DNA (nsDNA) through clamp opening at the rate *U*, returning to its apo form (magenta dot). All forms of apoParB can bind and unbind to nsDNA with rates *k*_on_ and *k*_off_, resp.

The LG is the simplest microscopic model for particles displaying short range interactions. It was historically derived from the Ising model, which, according to the mean-field solution [33], is known to exhibit a first-order phase transition in the presence of a non-zero magnetic field (of equivalently by favoring either holes or particle with a chemical potential in the LG formulation). The lattice framework is particularly adapted for modeling protein dynamics on DNA, as proteins rapidly move between discrete DNA binding sites. Notably, this class of LG models has namely been introduced to provide a microscopic foundation for hydrodynamic equations, such as the Navier-Stokes equations [34]. We define the LG model, previously proposed to describe the liquid-like behavior of ParB condensates [10]. The Hamiltonian of the LG model is:

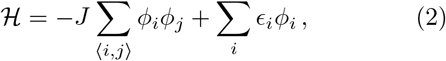

where *J* represents the contact interaction energy, *ϕ*_*i*_ is the occupation variable of the lattice site *i* (*ϕ*_*i*_ =0 or 1 if the site is empty or occupied, respectively) and ⟨*i, j*⟩ denotes all pairs of nearest neighbor sites on the lattice. The interaction energy *ϵ*_*i*_ of the site *i* is defined by *ϵ*_*i*_ = *ϵ*_s_ − *ϵ*_ns_, the difference between the affinity of ParB on specific DNA at *parS* (*ϵ*_s_) and non-specific DNA (nsDNA, *ϵ*_ns_). The lattice serves as a mean field representation of DNA, where each lattice site represents a potential 16bp ParB binding site [35]. Here, DNA refers to both chromosomal and plasmid DNA, present stochastically in the vicinity of *parS* since the plasmid is embedded in the nucleoid [36]. Thus, all ParBs are bound to DNA: either to *parS*, to *parS*-proximal DNA or to non-specific DNA, ı.e., the nucleoid.

Fig. 1B, presents the LG phase diagram in the (1*/*(*βJ*), *ρ*) plane where *β* = 1*/*(*kT*) is the inverse temperature, *k* the Boltzmann constant, *T* the absolute temperature and *ρ* the density. It is obtained in the mean field approach [32], yet it is a reasonable approximation to sketch the phenomenology of the realistic three dimensional LG. In this case, no exact solution exists and it requires thus to be studied numerically. At low interaction energy, *J*_V_, the system remains in a vapor-like state (green), unable to form droplets. For sufficiently high interaction energy *J*_LV_, the system is set in the LV phase (purple) below the coexistence line (full black line). At the coexistence line, the LG displays a first order (or discontinuous) transition. The LV phase is characterized by the existence of liquid droplet formation and consists of two regions: the metastable region (white and magenta stripes), located between the coexistence and the spinodal line (black dashed line) and the stable region, below the spinodal line. The difference between these two regions lies in kinetic effects. Indeed, the free energy minima defines the thermodynamic state of the system. In the metastable region, the free energy landscape exhibits a local minimum corresponding to a vapor-like state, separated by a barrier from the global minimum in the liquid–vapor coexistence phase. Thus in the metastable region, if the system is initially set in a disordered configuration, it must overcome the free energy barrier by thermal activation in order to switch to the minima in the liquid-vapor phase. Depending on the free energy barrier, the system may remain in the metastable vapor state for a prolonged period. For completeness, we notice the existence of a critical point at *ρ* = 1*/*2 and *T* = *T*_*c*_ (*T*_*c*_ = 3*/*2 in the mean field case), at which LG model undergoes a continuous (or second-order) phase transition.

The CS model, as depicted in Fig. 1C, involves the specific binding of apoParB (green dots) to *parS* sites (black square) at the binding rate *B*. This step is not rate-limiting due to the high affinity of ParB for *parS* (in the nanomolar range [6]) and its higher rate compared to other steps, as recapitulated in Ref. [13]. Binding to *parS* increases the affinity of ParB for CTP [17, 18]. Upon CTP binding, ParB undergoes a conformational change, forming a clamp-like structure (magenta circle) that sterically clashes with the DNA double helix, leading to ParB release from *parS* [9, 24]. *The combined rates of CTP binding and subsequent ParB release are modeled with a single rate R* (release rate), as the latter is the limiting step [13]. Once released, the clamped-ParB diffuses along the DNA (grey line) at the diffusion rate *D*. Lastly, upon CTP release, either through CTP unloading or CTP hydrolysis, ParB returns to its apo-form (magenta dots), which triggers unclamping and subsequent unloading from nsDNA at a rate *U*.

The novelty and main hypothesis of the present study is that ParB undergoes an activation switch from an inactive (green) to an active (magenta) form, with enhanced ParB-ParB interaction energy transitioning from *J*_V_ to *J*_LV_ (activated), where *J*_LV_ *> J*_V_. We propose that this switch occurs concomitantly with the release of ParB from *parS* in its clamp conformation. Note that both forms of ParB can bind to nsDNA, independently of the CS mechanism, with respective binding and unbinding rates *k*_on_ and *k*_off_.

The following section investigates numerically the type of activation necessary to trigger the phase transition in a biologically relevant timescale. To this end, we assay ParB macroscopic behavior with MC simulations and analyze the kinetics of droplet formation under activation in the LV phase.

### ParB droplet formation requires an activation of ParB-ParB interactions

According to the LG model, droplet formation is restricted to the LV phase. The question remains whether activation occurs within a biologically relevant timescale. To address this, we conducted Monte Carlo simulations [26] to study the kinetics of droplet formation. Our simulations were performed on a cubic lattice of size 160 × 40 × 40, where the lattice unit corresponds to the size of a ParB binding site, 16 bp of DNA [14–16]. The simulations were performed in the canonical ensemble, maintaining a fixed number of particles *N*_*b*_ to approximate the regulated number of intracellular ParB proteins (here *N*_*b*_ = 300). Note that the density is low (*ρ* ∼ 10^*−*4^) and we expect thus a first order transition, see Fig. 1B. The order parameter *ψ* is defined as the average density of bonds per particle:

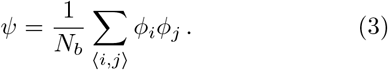

It distinguishes the vapor phase (*ψ* ≈ 0) from the LV phase (*ψ* ≈ *q*), where *q* is the coordination number of the lattice (here *q* = 6 for the simple cubic lattice). We started to establish the equilibrium phase diagram of the LG, shown in Fig. 2A, corresponding to biological parameters. The order parameter *ψ* is plotted as a function of the interaction energy *J*. Note that *J* is expressed in unit of *kT*. The red (increasing *J*) and blue (decreasing *J*) curves exhibit a hysteresis cycle, characteristic of a first-order phase transition. This defines three regimes: vapor (*J <* 2.7), metastable LV (2.7 *< J <* 3.1), and stable LV (*J >* 3.1). In brief, *J* was incremented (red curve) or decreased (blue curve) in discrete steps (Δ*J* = 0.06, see Mat. & Meth. for more details). We observe a first order transition at biological values of *J* ≈ 3. An activation from the vapor to the LV phase would correspond to a reasonable increase of *J*, by an increment of a few times the thermal energy.

**Figure 2.**
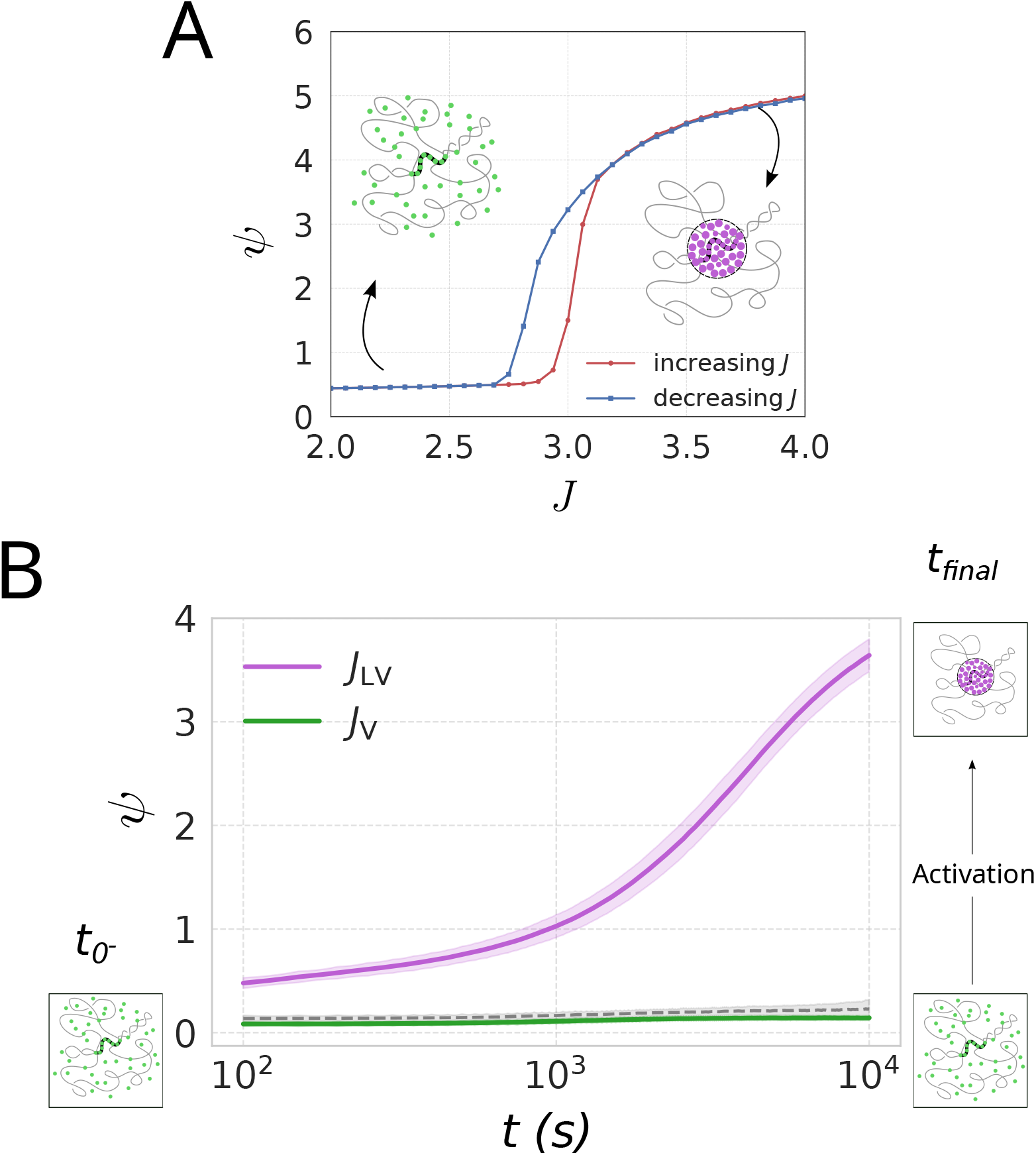
Partition complex formation needs an activation in the LV phase. **A** Equilibrium order parameter *ψ* as a function of the coupling constant *J* (expressed in unit of *kT*). The system, simulated on a 160×40×40 lattice with *N*_*b*_ = 300 particles. The red and blue curves represent increasing and decreasing *J*, respectively. At low and high values of *ψ*, the system is in the vapor and liquid-vapor (LV) phases, respectively. Illustrations representing the system’s state in each phase, using the same representation as Fig. 1. **B** Out-of-equilibrium order parameter versus time (same parameters as in panel A). The system, initially prepared in the vapor phase, relaxes towards equilibrium under two different conditions: activated (stable) LV phase (purple, *J* = 4.7) and vapor phase (green, *J* = 2.5). As a control, we plot in dashed grey an activation in the metastable region (*J* = 3).

In Fig. 2B, we plot the time evolution of *ψ* in order to study the out-of-equilibrium behavior of the droplet formation. Our protocol consists in preparing the system in a completely disordered state (vapor phase with *J* = 0) and let it evolve after activation in the LV phase (*J* = 4.7). This can also be interpreted as a quench, as increasing the coupling theoretically corresponds to decreasing the temperature. Under the experimental conditions used in this study, the cell cycle is approximately 60 minutes or 3.6 × 10^3^s, see [8]. Since one MC iteration approximates one second, the total simulation time of 10^4^s corresponds to about ten times the cell cycle. The purple curve describes the evolution of *ψ* when the LG is activated into the LV phase (*J* = 4.7). The droplet formation (defined as the time when *ψ* reaches half of its equilibrium value) occurs in average in 3 × 10^3^s corresponding to ≈45 minutes, based on 10^3^ simulations. Despite the simplicity of our approach and the most unfavorable initial conditions (ParB randomly distributed inside the cell instead of being clustered near *parS* because of the ParB operon close to *parS*), droplet formation occurs within a timescale corresponding to a fraction of the cell cycle. The green curve describes the evolution of *ψ* when the LG remains in the vapor phase (*J* = 2.5). No complex formation is observed, as it is the case in the absence of *parS* or certain ParB variants. As a control simulation, the gray dashed curve represents a quench into the metastable region (*J* = 3). Notably, green and gray curves are indistinguishable, with no nucleation observed within ten times the cell cycle, in agreement with Ref. [10]. Indeed, as described in Fig. 1, between the spinodal and the coexistence lines, the free energy exhibits a metastable vapor state in which the system can become trapped if the system is initially set in a disordered configuration. In Ref. [10], it was shown that nucleation triggered by ParB specifically bound to *parS* accelerates the transition by locally enhancing ParB-ParB interactions. However, this effect alone is insufficient to achieve droplet formation on a timescale compatible with the cell cycle. This supports that neither an activation of the interaction energy nor a fixed interaction within the metastable region is sufficient for complex formation.

These simulations support the hypothesis that ParB must undergo an activation of its interaction energy to reproduce the observed experimental phenomenology: (i) the absence of complex formation with Δ*parS* or some ParB variants, and (ii) the formation of a complex within the timescale of a fraction of the cell cycle with a functional ParAB*S* system. In the following, we present experimental evidence demonstrating that a complete ParB-CTP cycle is required for this phase transition and subsequent droplet formation.

### A functional ParB-CTP cycle is essential for droplet formation

To support our modeling, we show here that altering the release of ParB_F_ from *parS*_F_ or impairing the unloading of clamped ParB_F_ hinders activation and consequently prevents droplet formation. To test this, we investigated the ParAB*S*_F_ system of plasmid F, specifically targeting mutations in the boxIII motif of the *parB*_F_ gene which, along with the boxII motif, is involved in CTP binding and hydrolysis [17, 18]. The boxIII motif contains highly conserved asparagine (N) and arginine (R) residues (Fig. 2A). We generated two ParB_F_ variants by substituting these residues with alanine (A), resulting in ParB_F_-N153A and ParB_F_-R156A. The resulting mutations, ParB_F_-N153A and ParB_F_-R156A, strongly impair the stability of mini-F plasmids, with high loss rates compared to parB wt (Table S1).

To characterize *in vivo* at which stage these ParB variants are deficient in the partition complex assembly, we started by analyzing the expression levels of these variants on mini-F plasmids (Fig. S1). ParB_F_-R156A and ParB_F_-N153A were expressed at 1.2- and 3.4-fold higher levels, respectively, compared to ParB WT, respectively. Since ParA_F_ acts as the auto-repressor of the *par* operon, which repressor activity is stimulated by 20- to 40-fold through interaction with ParB_F_ and further enhanced 3- to 4-fold in the presence of *parS*_F_ [23, 38], these results suggest that the ParA_F_-ParB_F_ interactions remain fully functional for both variants. However, the lack of additional repression in the presence of *parS*_F_ for ParB_F_-N153A indicates that this variant is specifically defective in *parS*_F_ -mediated stimulation.

Next, using fluorescence microscopy, we investigated partition complex assembly. WT ParB_F_-mVenus formed bright and distinct foci in the presence of *parS*_F_ but not in the absence of *parS*_F_ (compare Fig. 3B(a) with 3B(b) [5]). The N153A variant showed diffuse fluorescence with no foci formation (Fig. 3B(c)), indicating a failure to assemble partition complexes. The R156A variant displayed an intermediate phenotype, with reduced foci intensity and increased diffuse fluorescence (Fig. 3B(d)), suggesting partial defects in partition complex assembly.

**Figure 3.**
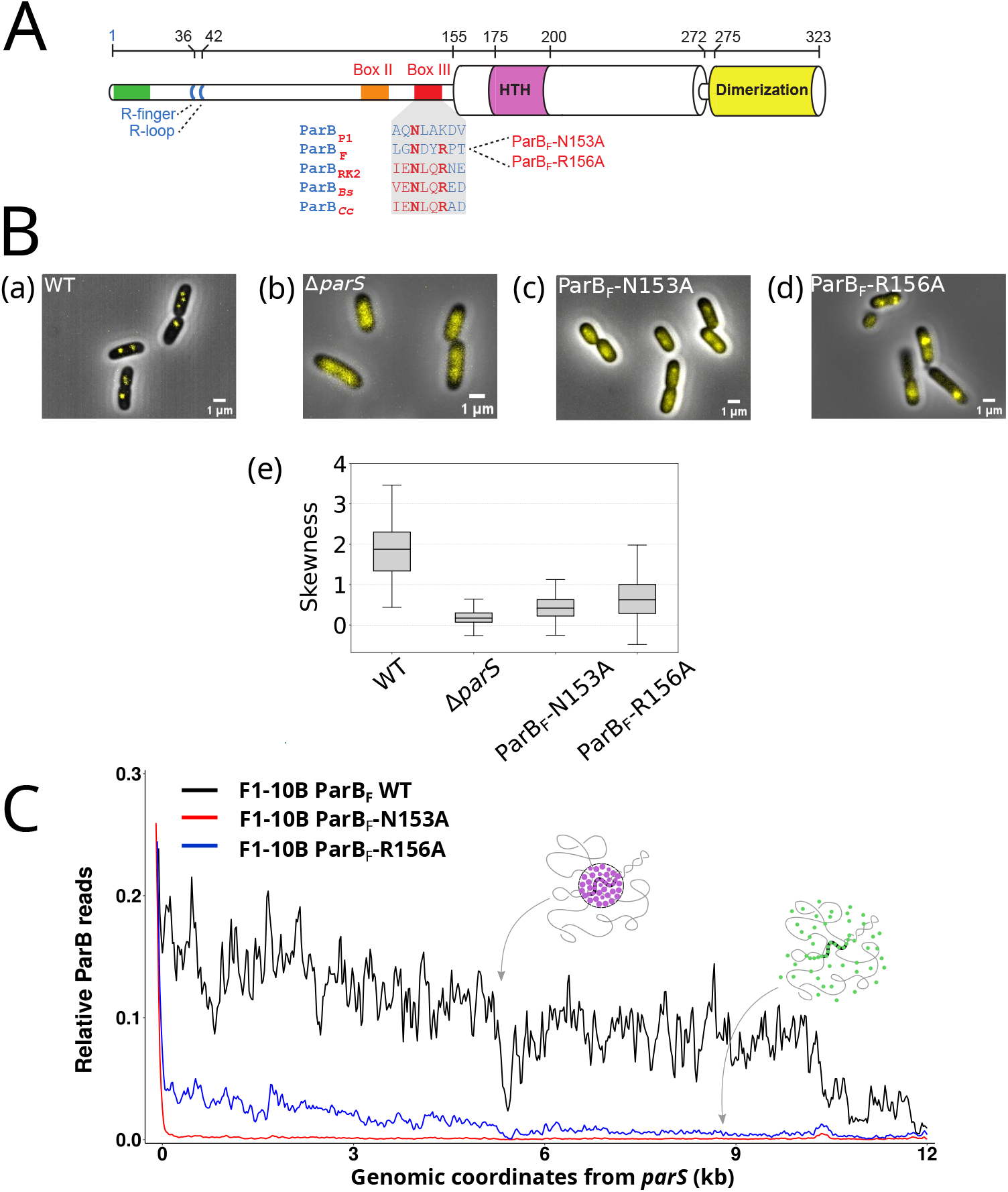
The ParB-CTP cycle is essential for ParB droplet formation. **A** Schematic representation of ParB_F_. (Top) Amino acid (aa) numbering to scale. (Middle) Functional domains (Helix-Turn-Helix and dimerization) and motifs (BoxII and III) are depicted. Thick cylinders (from aa 155–272 and 275–323) represent regions of ParB_F_ resolved by X-ray crystallography, while thin cylinders (from aa 1–155 and 272–275) correspond to unresolved regions [37]. (Bottom) Conservation of the aa sequence in the BoxIII. Conserved residues from plasmids P1, F and RK2 and from the chromosomes of *Bacillus subtilis* (*Bs*) and *Caulobacter crescentus* (*Ccre*) are colored in red. **B** Fluorescence imaging of ParB_F_ fused with mVenus fluorescent tag *in vivo. E. coli* cells, grown in exponential phase, exhibit ParB_F_-mVenus fluorescence expressed from the endogenous genetic locus of miniF plasmids. Representative images (a)-(d) and skewness analysis (e) of cells with miniF carying a deletion (Δ*parS*) or not of *parS* and expressing ParB_F_ WT or the variants ParB_F_-N153A ParB_F_-R156A. Scale bars: 1 *µm*. **C** The normalized average ParB reads are shown as a function of genomic coordinates from the right of *parS* over a 12.5 kb region for ParB_F_ WT (black), ParB_F_-N153A (green), and ParB_F_-R156A (blue) variants. These profiles were determined by ChIP-sequencing from biological duplicates using *E. coli* cells carrying F1-10B derivatives. Additionally, an illustration of the resulting ParB macroscopic behavior from the model is provided, corresponding to Fig. 1A. Detailed ParB read distributions for each replicate, along with input data, are provided in Fig. S2-S4.

To further analyze the assembly defects, we performed ChIP-sequencing (ChIP-seq; Table S2), which quantifies ParB enrichment along the plasmid DNA [5, 8]. Enrichment at and near *parS* sites serves as a proxy for ParB droplet formation. In the absence of droplet formation, ParB enrichment is restricted to *parS* or its close vicinity, with a homogeneous non-specific distribution of ParB across the whole genome generates background noise in ChIP-seq profiles [8]. We focused on the right-hand side relative to *parS*, as enrichment decay spans larger genomic range, while the left-hand side is constrained by a roadblock as previously reported (Fig. S2) [5]. In Fig. 3C, the WT profile (black curve) shows ParB enrichment peaking at *parS*, followed by a rapid drop and long-range decay over ∼15 kb. As previous studied [5, 8, 39], this decay is described by a power law obtained with the stochastic binding model, where ParB forms a droplet centered on *parS*. After normalization, the integral of the profile indicates ∼ 120 ParB molecules in the enriched region, consistent with the estimated ∼ 250 ParB dimers per complex [5] (with the remainder of ParB being likely bound to the chromosome). For the N153A variant (red curve and Fig. S3), enrichment is observed exclusively at *parS*. Beyond *parS*, the profile immediately drops to basal levels. The total ParB count, derived from the profile integral, is ∼10, primarily reflecting *parS*-bound ParB. This indicates that while ParB_F_-N153A is fully proficient in *parS* binding, it is deficient in undergoing the CTP-induced conformational change necessary to transition into the clamped form and recruit additional ParB molecules. Consequently, this variant is unable to convert into the activated form required for droplet formation. In contrast, the R156A variant (blue curve, and Fig. S4) shows the characteristic drop after *parS* and a progressive binding decay over 12 kb. The integral of this profile corresponds to ∼30 ParB molecules. This decay likely arises from both *parS*-bound ParB and sliding clamps, consistent with the CS model where the release and unloading parameter remain unchanged (Fig. 4 and analyses below), and thus indicates that ParB_F_-R156A is proficient for binding to *parS* and CTP.

**Figure 4.**
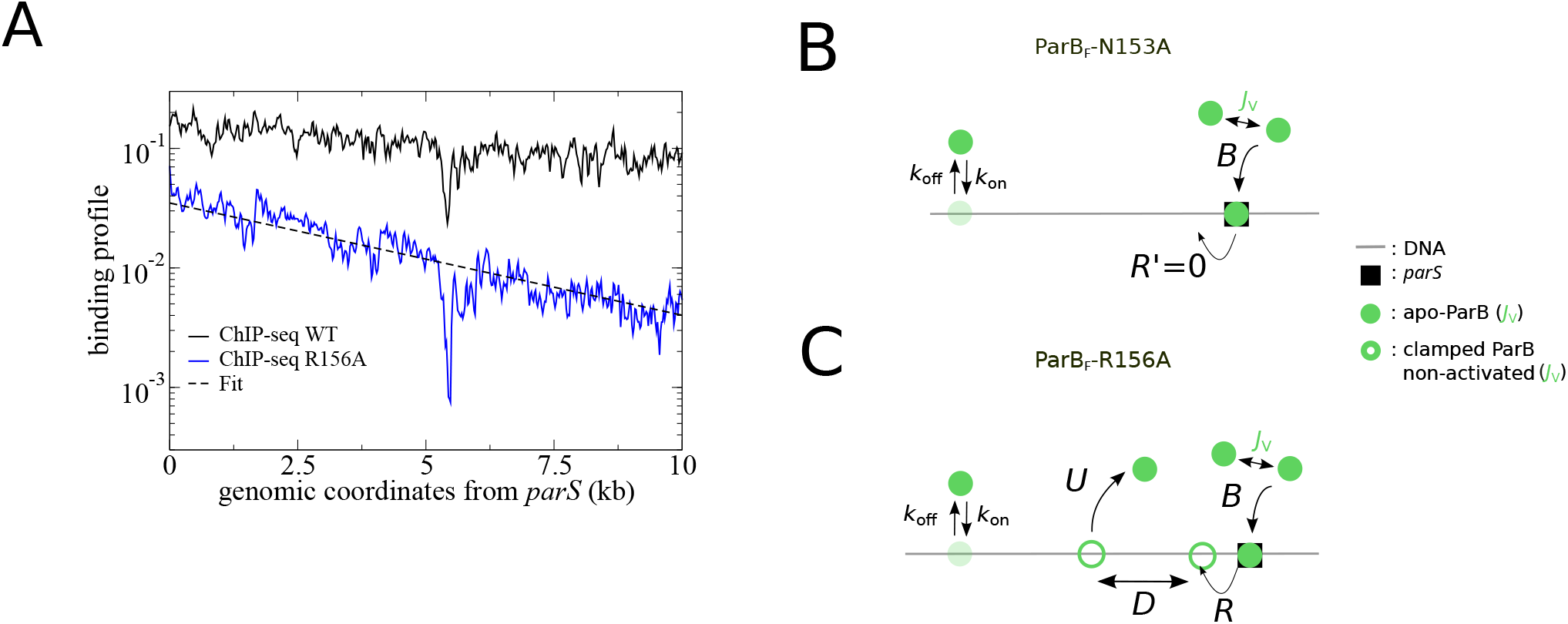
**A** The ChIP-seq ParB profile of the ParB_F_-R156A variant corresponds to the C&S model alone. ParB DNA binding profile for ParB WT (black line) and for ParB_F_-R156A (blue line) are plotted versus the genomic coordinate from *parS*_F_ in a log-lin scale. The black dashed line corresponds to the fit according to the C&S model with parameters close to the WT (see text for details). **B** ParB*-N153A variant is blocked in a parS*_*F*_ *-bound form (R*^*′*^ = 0). ParB-N153A (green dots) specifically binds to *parS* (black square) at the rate *B*. By contrast to ParB WT, it remains bounds to *parS* without transitioning into the clamped state. As a result, ParB cannot diffuse along the DNA. ParB may bind and unbind to non-specific DNA (nsDNA) with rates *k*_on_ and *k*_off_, respectively, but do not undergo a transition to the activated form for enhanced ParB-ParB interactions. **C** *ParB-R156A variant is unable to promote enhanced ParB-ParB interactions*. ParB-R156A (green dots) binds to *parS* at the same rate *B*. The clamp is released from *parS*, diffuses along DNA and unloads at rates *R, D* and *U*, respectively. The unloading from nsDNA at rate *U* remains unchanged. By contrast to ParB WT, ParB-R156A clamps are not activated for enhanced ParB-ParB interactions, preventing the formation of liquid-like ParB droplets via phase transition.

While this mutation permits the formation and diffusion of ParB clamps, it likely impairs the recruitment of additional ParB molecules, as supported by the reduced number of bound molecules in ChIP-seq. We cannot exclude, however, that the kinetics of clamp formation or DNA unloading are also affected, potentially contributing to the narrower DNA occupancy profile observed for the mutant. This data thus strongly suggests that this variant does not transition to an activated form needed for phase transition.

To emphasize the exponential decay of the ParB_F_-R156A profile, the data were plotted in a log-lin scale (Fig. 4A). According to the CS model [13], the profile is fitted by an exponential as *y* = *a* exp(− *bx*) (black dashed line), where *a* = 0.035 and *b* = 2.162 × 10^*−*4^bp^*−*1^. The characteristic length of the exponential is thus 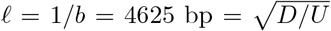. Using the value *D* = 0.05*µm*^2^ s^*−*1^ [10], which is in agreement with the one of ParB_*Mxan*_ in *M. Xanthus* (0.02*µm*^2^ s^*−*1^; [9]), it comes *U* = *D/*𝓁^2^ = 0.02s^*−*1^ in very good agreement with the value *U*_*Ccre*_ = 0.017*s*^*−*1^ obtained with ParB_*Ccre*_ [19]. The amplitude of the profile 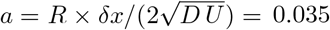 leads to *R* = 0.37s^*−*1^, larger by a factor ∼4 than the *in vitro* value *R*_*Ccre*_ = 0.1s^*−*1^ in [19] but still of the same order of magnitude. Overall, the kinetic parameters of the variant R156A are largely comparable to those of ParB WT. We translate these findings into the mechanistic framework of the CS model. For N153A (Fig. 4B), ChIP-seq and microscopy reveal no enrichment on nsDNA but only at *parS*, with no visible ParB droplets. Compared to the WT model (described in Fig. 1C), this corresponds to a new set of parameters where *parS* binding occurs normally at the rate *B*, but a modified release rate *R*^*′*^ = 0 prevents the clamped form and activation switch, thereby inhibiting droplet formation. For ParB-R156A (Fig. 4C), ChIP-seq indicates residual enrichment consistent with CS (as modeled in Fig. 4A) but not with droplet formation, supported by microscopy. This latter mutation likely impairs the activation switch required for ParB droplets formation but leaves other parameters unchanged.

These findings underscore the necessity of a fully functional ParB-CTP cycle for droplet formation, highlighting the role of ParB clamp activation in this process. The mutations studied here reveal distinct effects on ParB dynamics, demonstrating how disruption of either CTP binding and/or hydrolysis impairs LLPS and partition complex assembly.

## Discussion

The partition complex exhibits liquid-like properties [10, 11], yet the mechanisms controlling ParB droplet formation remain incompletely understood. While the necessity of a functional *parS* site for droplet formation is well established [5, 8], the precise role of CTP in this process remains unclear. Recent studies have proposed the Clamping & Sliding (CS) model for partition complex assembly, which outlines a ParB-CTP cycle involving specific ParB binding to *parS*, followed by CTP binding, clamp formation, *parS*-release and CTP release-dependent hydrolysis DNA unloading [13]. However, this model accounts for only a small fraction of ParB molecules (*<*10) bound to DNA, significantly fewer than the estimated ∼ 250 ParB molecules per partition complex [5]. Moreover, the CS model does not inherently rely on the liquid-like properties observed in ParB condensates. These discrepancies highlight the need to identify additional mechanisms, such as a CTP-dependent activation step of the ParB-ParB interactions during the CS cycle, to fully explain droplet formation. Reconciling the CS model with the liquid-like behavior of ParB droplets is therefore essential for understanding the assembly and function of the partition complex.

In this study, we propose that ParB activation for liquid-like behavior coincides with its transition into the clamped conformation. Evidence suggests that the synergistic effect of *parS* and CTP binding allows ParB to adopt a distinct conformational state [9, 18]. We infer that these *parS*-activated ParB-CTP forms persist after *parS*_F_ releases and enhance ParB_F_-ParB_F_ interactions. These two populations, the clamped-form and, after unloading, the open-ParB form are structurally distinct, especially in their N-terminal domain [17, 18]. Although the strength of ParB–ParB interactions is likely to differ between these two forms, we speculate that the open ParB form may transiently retain an activated state that promotes enhanced ParB-ParB interactions after release. In spite of the fact that an increase in ParB-ParB interaction strength remains to be formally demonstrated, magnetic tweezers experiments (refs [40, 41]) have shown that CTP promotes multiple ParB_F_ loading onto *parS*_F_-containing DNA. This loading leads to the formation of condensed partition complex-like assemblies, also strongly suggesting that CTP-bound ParB_F_ adopts a conformation that enhances ParB-ParB interactions. Moreover, the recently shown capability of clamped-ParB to recruit apo-ParB and convert them, independently of *parS*, in a clamped form able to diffuse on an adjacent DNA (*i*.*e*., *trans* loading; [21]) is in agreement with our model. Indeed, such conversion is blocked in the presence of CTP*γ*S by preventing the switch to an activated form, and thus the formation of transient ParB-ParB bridge. We propose that this activation step serves as a critical switch, ensuring that ParB droplets form exclusively at *parS* sites, thereby preventing nonspecific aggregation elsewhere in the genome. Such precise localization is essential for maintaining cellular functionality and ensuring efficient DNA partitioning during the cell cycle. Our hypothesis is supported by both simulations and experimental evidence, which collectively highlight the central role of this activation step in coupling ParB droplet formation with its functional specificity.

To decipher the mechanisms of droplets formation, kinetic simulations using the LG model were employed to assess the timescale of this process. Knowing that droplets can only form in the LV phase, we initialized the system in the vapor phase and activated it to transition into the LV phase. Our simulations revealed that droplet formation occurs within a timescale corresponding to a fraction of the cell cycle. Our findings indicates that ParB cannot maintain a constant interaction energy (in the metastable region) to account for a droplet formation in a timescale compatible with the cell cycle even with the presence of a nucleation seed provided by *parS*, as previously proposed [10]. Instead, an additional mechanism is required to enhance ParB-ParB interactions in the presence of *parS*. Our experimental results, using ParB variants of the CTP binding pocket, support this conclusion. Although boxes II and III are highly conserved, the mutation of a conserved residue does not necessarily lead to the same change in ParB activity. For instance, mutation of the highly conserved Asn residue in boxIII (Fig. 3A) leads to a three-fold decrease in CTP-binding affinity for ParB of *M. xanthus* [9], whereas it does not impair CTP binding for ParB of *B. subtilis* [18]; nonetheless, both variants are deficient in CTP hydrolysis and clamp formation [17, 18]. For ParB of plasmid F, the N153A mutation prevents ParB_F_ to be released from *parS*_F_ as a sliding clamp (Fig. 3C and S3), behaving as a CTP-binding deficient variant that is unable to form ParB droplets. Such deficiency was also observed for ParB_F_ mutation in boxII (ParB_F_-3R [8]). This indicates that the CTP-dependent conversion to a clamp form is necessary to enable phase transition. However, this conversion is not sufficient for droplet formation, as indicated by the ParB_F_-R156A variant, which binds *parS*_F_ and exhibits the CTP-dependent *parS*_F_ release (Fig. 3C and Fig. S4). After release, it diffuses from *parS*_F_ as sliding clamp since its DNA profile corresponds almost perfectly with the prediction of the CS mechanism alone (Fig. 4A). ParB_F_-R156A variant does not transition to recruit additional ParB, lacking the activation of ParB-ParB interactions. This indicates that the clamped form of ParB_F_-R156A is locked in a conformation that prevents the transition required for phase separation. These results can be interpreted within the framework of the CS model as follows: (i) ParB-N153A represents a system with no clamped-ParB formation and thus no activation (*R*^*′*^ = 0), and ParB-R156A represents a clamped form locked in an inactive conformation. In both cases, the failure of ParB to switch to an activated form with enhanced ParB_F_-ParB_F_ interactions prevents droplet formation and partition complex assembly. These findings underscore the critical requirement not only for a fully functional CS cycle, but also for a finely tuned activation step during the ParB-CTP cycle to drive droplet formation and ensure efficient DNA partitioning.

To go beyond the lattice model and account for DNA fluctuations, the LG must be embedded on a fluctuating polymer. This system falls within the universality class of magnetic polymers [42, 43]. Polymer fluctuations are expected to smoothen the transition. Magnetic polymers are known to exhibit a weak first-order transition that becomes continuous when the applied field is sufficiently strong. Further studies will be required to estimate the relevant field strength in relation to our experimental concentrations. Additionally, to better mimic the intracellular environment, the system should be embedded in a melt of magnetic polymers, reflecting the effect of the nucleoid.

In conclusion, we propose that CTP functions as a molecular switch, driving ParB from a vapor to a LV phase. The activation performed by coupling phase separation to ParB-CTP conformational changes ensures both the specificity and efficiency of ParB droplet formation at *parS*, preventing nonspecific aggregation elsewhere in the genome. This represents the first identified CTP-controlled switching mechanism regulating liquid-like droplet formation. Further investigations are needed to determine whether this switch can be directly triggered through ParB-ParB interactions upon molecular contact. Understanding these interactions could provide broader insights into how CTP modulates phase transitions in cellular processes and partitioning mechanisms.

## Financial disclosure

This work was supported by the CNRS 80Prime MITI grant (ANCODS) and by the Agence National pour la Recherche (ANR-24-CE12-1319).

## Acknowledgements

We thank G. Morla for its early involvement in the project during his master’s studies. We acknowledge all members of the GeDy (CBI) and the SCPN (L2C) teams for fruitful discussions, and the staff at the GeT-Biopuces platform (INSA, Toulouse) for handling libraries and sequencing of ChIP-seq samples. This work was supported by the CNRS 80Prime MITI grant (AN-CODS) and the ANR (BaDS; ANR-24-CE12-1319-01).

## Data availability

The raw and processed ChIP-seq data reported in this paper have been deposited in Gene Expression Omnibus database (GEO, NCBI) and are accessible through GEO Series accession numbers GSE256357 (ParB_F_) and GSE289093 (ParB_F_-N153A and ParB_F_-N156A). The custom codes are available on the generic repository Zenodo with the permanent DOI 10.5281/zenodo.16882674

## Notes

### Competing Interest Statement

The authors have declared no competing interest.

### Summary of Updates

Minor changes after revision of peer referees

