## Supplementary Tables and Figures for "The ParB-CTP Cycle Activates Phase Separation in Bacterial DNA Segregation"

**Table S1.** *ParB variants in boxIII are impaired for partitioning activity in vivo.* The functionality of fluorescently-tagged proteins was assayed by measuring the plasmid loss per generation. Number of independent experiments is indicated in brackets. * Note that the data for the mini-F $\delta parS$ was performed with untagged ParB from our previous work [1].

| Mini-F Variant | Characteristics | Loss Rate (%) |
|---|---|---|
| pJYB234 | par A, B-mVenus, S | 0.08 (1) |
| pDAG115 | par A, B, ΔS | 7.90 (*) |
| pJYB341 | par A, B-N153A-mVenus, S | 7.42 (3) |
| pJYB342 | par A, B-R156A-mVenus, S | 2.40 (3) |

**Table S2:** Synoptic of the information relative to the ChIP-sequencing experiments

| Strain names | Main properties | Growth condition[a] | ChIP Input/IP Antibody* | Av. Library size** | Reads total | Reads mapped | Repli-cates *** |
|---|---|---|---|---|---|---|---|
| DLT4098 | E. coli / F1-10B (ParB WT) | 37°C | Input<br>IP anti $ParB_F$ | 151<br>162 | 5,543,092<br>18,110,058 | 5,518,379<br>18,000,450 | R1<br>R1 |
| DLT4115 | E. coli F1-10B16 (ParB-N153A) | 37°C | Input<br>IP anti $ParB_F$ | 182<br>173<br>181 | 4,425,661<br>15,852,048<br>12,271,455 | 4,412,379<br>1,5493,504<br>9,535,898 | R2<br>R1<br>R2 |
| DLT4116 | E. coli F1-10B17 (ParB-R156A) | 37°C | Input<br>IP anti $ParB_F$ | 162<br>164<br>194 | 6,580,179<br>22,526,673<br>14,083,917 | 6,470,513<br>21,906,102<br>13,693,419 | R1<br>R1<br>R2 |

* Affinity-purified antibodies from rabbit-serum.

** Average size of the DNA fragments in the library as measured by sequencing.

*** Replicates indicated in bold correspond to the ones presented in figures in the manuscript.

**Table S3:** *Bacterial strains and plasmids.* All *Escherichia coli* strains are derivatives of *E. coli* K12.

| Strains | Genotype/relevant properties | Source/references |
|---|---|---|
| DLT1215 | *F⁻, thi leu thyA deoB supE rpsL, Δ(ara-leu, zac3051::Tn10* | (Bouet et al, 2005) |
| DLT1780 | DLT1215 / pDAG114 | (Sanchez et al, 2015) |
| DLT3053 | DLT1215 *Hu-mCherry*, FRT-Kan-FRT | (Le Gall et al, 2016) |
| DLT3055 | DLT3053 / pJYB234 | (Debaugny et al., 2018) |
| DLT3586 | DLT1215 / F1-10B | (Debaugny et al., 2018) |
| DLT3589 | DLT1215 / F1-10B *parB*-mVenus | (Debaugny et al., 2018) |
| DLT4050 | DLT3053 / pJYB341 | This work |
| DLT4051 | DLT3053 / pJYB342 | This work |
| DLT4052 | DLT3053 / pJYB343 | This work |
| DLT4053 | DLT3053 / pJYB344 | This work |
| DLT4113 | DLT1215 / F1-10B14 | This work |
| DLT4114 | DLT1215 / F1-10B15 | This work |
| DLT4115 | DLT1215 / F1-10B16 | This work |
| DLT4116 | DLT1215 / F1-10B17 | This work |

| Plasmids | Relevant characteristics | Source/references |
|---|---|---|
| F1-10B | $F^+$, *lac*$^+$ tn10, *ccdB*$^-$ *cat*$^+$ | (Debaugny et al., 2018) |
| F1-10B-BmV | F1-10B, *parB*$_F$*-mVenus* | (Debaugny et al., 2018) |
| F1-10B14 | F1-10B-BmV *parB*$_F$-N153A | This work |
| F1-10B15 | F1-10B-BmV *parB*$_F$-R156A | This work |
| F1-10B16 | F1-10B *parB*$_F$-N153A | This work |
| F1-10B17 | F1-10B *parB*$_F$-R156A | This work |
| pDAG114 | mini-F, *repFIA*$^+$, *ccdB*$^-$, *resD*$^+$, *rsfF*$^+$, *cat*$^+$ | (Lemonnier et al, 2000) |
| pDAG115 | pDAG114 Δ*parS*$_F$ | (Lemonnier et al, 2000) |
| pJYB234 | pDAG114, *parB*$_F$*-mVenus* | (Debaugny et al., 2018) |
| pJYB341 | pJYB234, *parB*$_F$-N153A-*mVenus* | This work |
| pJYB342 | pJYB234, *parB*$_F$-R156A-*mVenus* | This work |
| pJYB343 | pDAG114, *parB*$_F$-N153A | This work |
| pJYB344 | pDAG114, *parB*$_F$-R156A | This work |

### Supplementary Figures

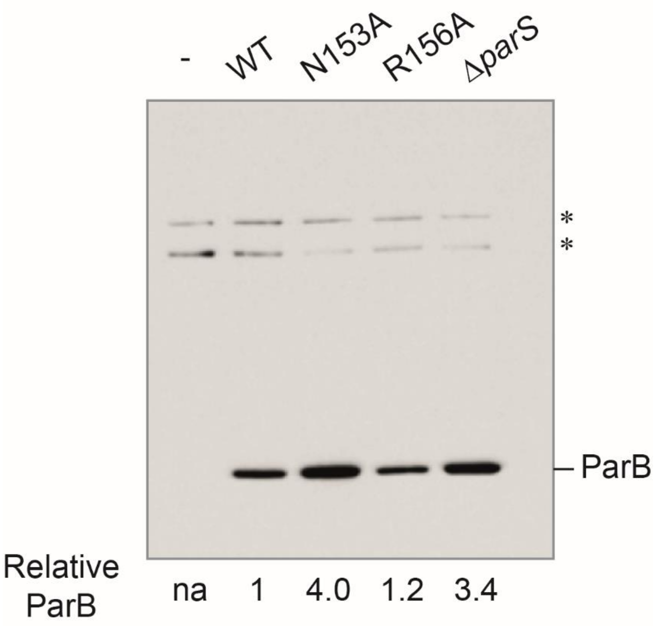


**Figure S1.** *Expression level of ParB variants.* Cultures of *E. coli* strains carrying or not (-) mini-F plasmids encoding *parB* WT, *parB*-N153A, *parB*-R156A or deleted from *parS* were grown in exponential phase. Samples were harvested and protein extracts were resolved on a 4-12% Bis-Tris gradient gel (Invitrogen). A typical Western blot using anti-ParB antibodies is displayed for DLT1215 strains, carrying or not (-) pDAG114 derivatives with the indicated *parB* alleles, grown in exponential phase the indicated plasmids. The level of ParB relative to WT ParB was quantified and indicated below. This relative value was normalized with the two non-specific cross-reactive bands (*) on top of the gels.

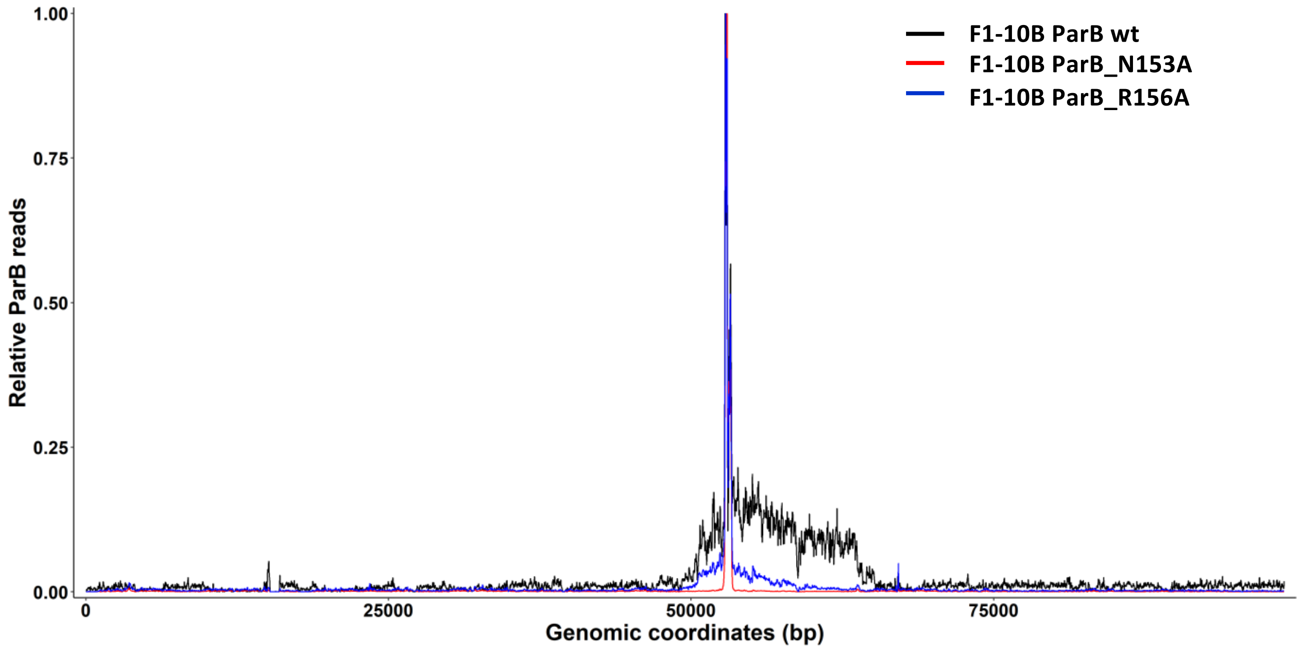


**Figure S2.** *ParB DNA binding profiles on plasmids F1-10B.* Density of the binned ParB reads from ChIP-seq of biological duplicates performed on plasmids F1-10B are displayed in black (ParB WT), red (ParB-N153A) and blue (ParB-R156A) over the F1-10B genome (same data as in Figures S3 and S4; 1 is the maximum density in a bin).

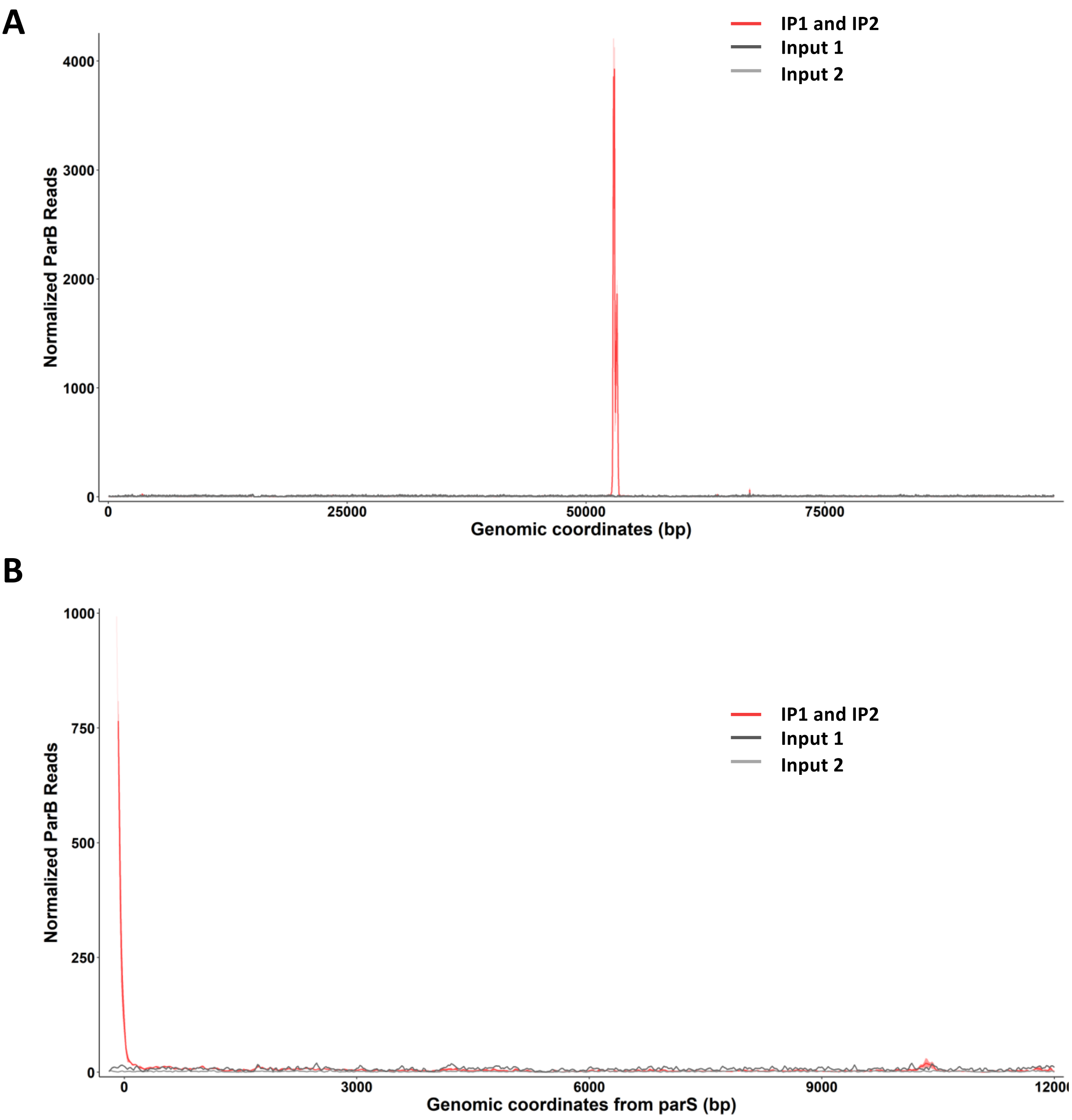


**Figure S3.** *ChIP-seq profile of ParB-N153A variant.* Normalized ParB reads are displayed (ribbon representation) as a function of the genomic coordinates of the plasmid F1-10B, with the red line representing the average of two biological replicates (IP1 and IP2). The differences between the duplicates are represented by the light red ribbon. Inputs correspond to the sequencing of DNA preparation before IP (dark and grey lines). **A** Normalized ParB-N153A reads over the whole genome of F1-10B. **B** Zoomed of the normalized ParB-N153A on the right side of $parS_F$ over 12.5-kbp. Coordinates are relative to the last bp of the last $parS$ site.

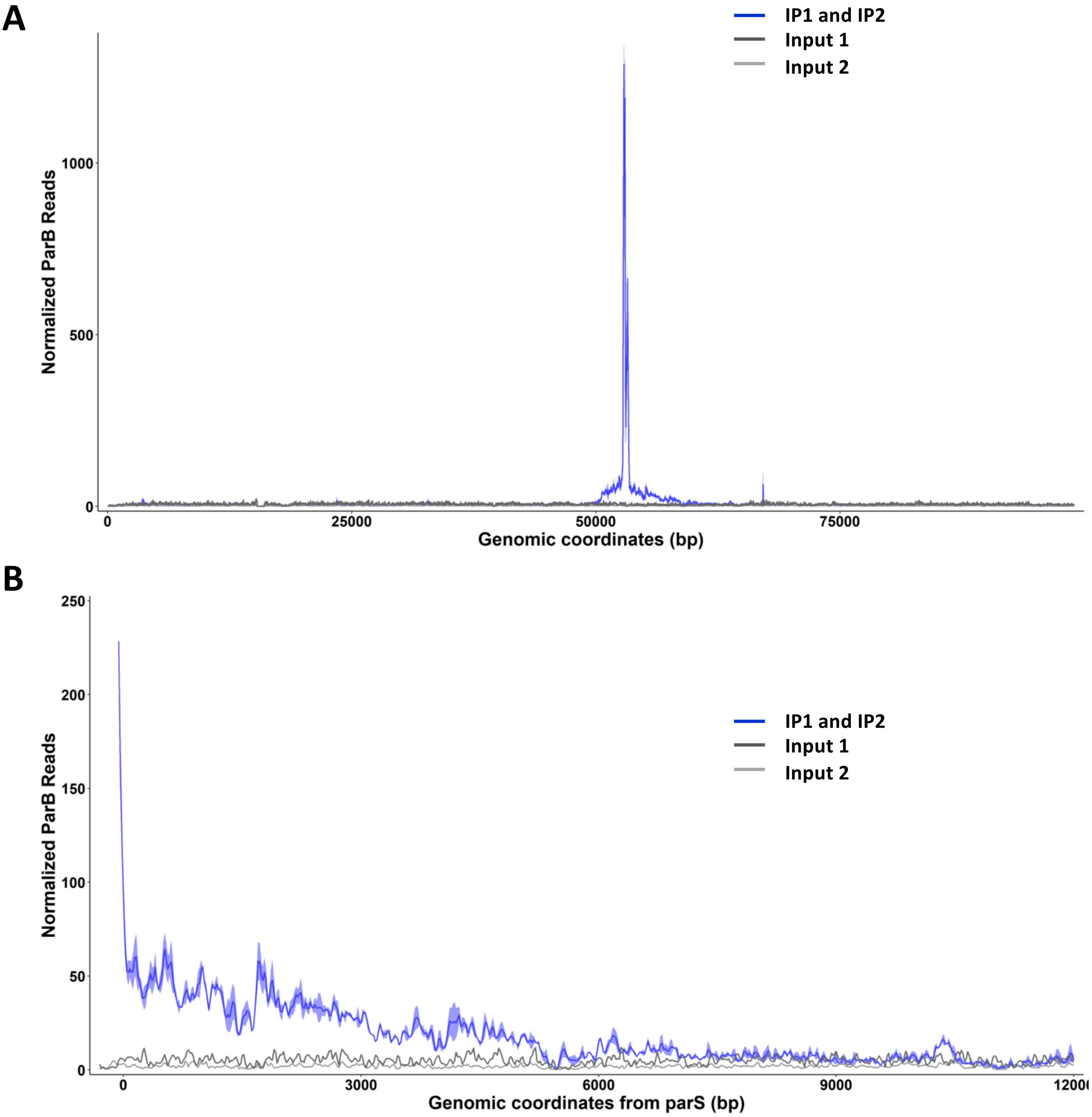


**Figure S4.** *ChIP-seq profil e of ParB-R156A variant.* Normalized ParB reads are displayed as in Figure S3 with the average of two biological replicated (IP1 and IP2) represented with a blue line. **A** Normalized ParB-R156A reads over the whole genome of F1-10B. **B** Zoomed of the normalized ParB-R156A on the right side of $parS_F$ over 12-kbp. Coordinates are relative to the last bp of the last *parS* site.
